# Sputum-spotted solid matrix designed to release diagnostic-grade *Mycobacterium tuberculosis* DNA demonstrate optimal biocontainment property

**DOI:** 10.1101/2023.08.25.554768

**Authors:** Krishna H. Goyani, Pratap N. Mukhopadhyaya

## Abstract

Biosafety quotient of a reagent-coated cellulose matrix designed to release *Mycobacterium tuberculosis* (Mtb) DNA from sputum deposited onto it, was determined. Thirty-seven sputum samples infected with *Mtb* and 5 sputum samples from healthy individuals were processed and spotted onto the TBSend cards. Live *Mtb* bacilli were attempted for rescue from the spotted TBSend cards by washing them in phosphate buffer under mild shaking conditions. No live *Mtb* bacilli could be detected by the BACTEC™ MGIT 960™ TB System from both *Mtb* infected as well as non-infected, sample-spotted TBSend cards when they were washed at two different time-points, *viz*., 1 minute and 15 hours from the time of spotting of the cards with sputum mixed with the preprocessing “spotting” buffer. The study reiterated published finding that chaotropic agents, which is an active component of the TBSend card module, have cent per cent bactericidal properties with regard to *Mtb*.

## Introduction

There has been a dramatic improvement in approach for addressing health threat posed by Tuberculosis (TB) disease following implementation of the Millennium Development goals which aimed to reduce to half, the worldwide mortality and morbidity caused due to TB between the period of 1990 and 2015. Despite this tremendous global effort, the TB continues to be a major cause of mortality and morbidity arising from any single infecting pathogen across the world (World Health Organization, 2015).

The transmission of this pathogen is dictated by the environment, type of host and several other factors. For infections that arise due to direct transmission of the pathogen, it is necessary to understand the potential for infection of the index patients in the disease zones (Woolhouse *et al*., 1997; Galvani and May, 2005]. The rate of infection from a single patient depends on individual characteristics, with some people infecting a larger number of uninfected persons while others, infecting a fewer number of them or sometimes, none (Wang *et al*., 2008). TB infection occurs when the bacilli come out from an infected patient and get dispersed in the air and eventually reach the alveoli of another host human. The defence mechanism of the host however phagocytizes the pathogen using its alveolar macrophages that constitute the innate immune defence mechanism of the host (Urdahl *et al*., 2011). However, this infection progresses towards disease when few of the microbes escape the innate defence and replicate actively inside the macrophages. They then migrate to other cells in the vicinity that include epithelial as well as endothelial cells and soon reach significantly high microbial burdens (Wolf *et al*., 2008).

The process of coughing by infected patients releases the highest quantum of droplets of various sizes (Yang *et al*., 2007). It is therefore imperative that great precautions are taken while handing sputum of patients suffering from TB as this is one of the most potent sources of spread of the disease in the population. (Bhatt *et al*., 2012; Jangid *et al*., *2016*).

*Mycobacteria* bacilli has the potential to retain viability for several months on dry surfaces. It has been demonstrated that *M. bovis* can survive on moisture-free surfaces at 4-degree C. Similarly, *Mtb* has been shown to survive in cockroach faeces for 8 weeks, on carpets for 19 days, on dry wood for more than 88 days, in wet as well as dry soil for over 4 weeks and in the environment in general for over 74 days, especially when the bacilli is protected from intense light sources (Kramer *et al*., 2006; Phillips *et al*., 2003).

The transportation process of sputum for diagnostic purposes has significant potential for spread of TB disease and is an underestimated cause of dissemination of the pathogen to people engaged in the process who eventually become potential secondary spreaders of the disease. An Indian study revealed that only around 33% of doctors in the private sector use proper and prescribed method of transportation of sputum for its testing (Basu *et al*., 2013). In another study it was found that only half of the patients surveyed followed appropriate sputum transportation practices (Rekha *et al*., 2013). Therefore, safe sputum disposal and transportation is a key factor for proper control of spread of this disease (World Health Organization, 2007) along with other activities to enhance awareness among common man and the medical fraternity, which all together play a crucial role in TB control strategies (TBC India, 2013).

In this study, we demonstrate biocontainment quotient of a solid-matrix-based sputum transportation, storage and DNA-release device called the TBSend card and its potential to address biosafety concerns when used for room temperature transportation, extended period of storage at non-refrigerated condition and brief processing to release diagnostic-grade DNA for Nucleic Acid Amplification Testing (NAAT) including its cartridge-based variants (CB-NAAT).

## Materials and Methods

### Collection of clinical samples

Thirty-seven smear-positive sputum samples from presumptive TB patients, screened from a total of 58 individuals were used in this study. All samples were collected from walk-in patients (3 ml) visiting the Microcare laboratory and Tuberculosis Research Center (Surat, Gujarat), a TB testing facility accredited by the Central TB Division, Ministry of Health, Government of India. Samples were collected between the period of 2017-2019 in four batches, namely batch 1, 2, 3 & 4 and comprising of 12, 18, 7 and 5 sputum samples respectively. Batch 1, 2 and 3 comprised of TB infected sputum samples while batch 4 comprised of sputum from healthy individuals (Figure 1). *Processing of sputum samples*

**Figure 1.**
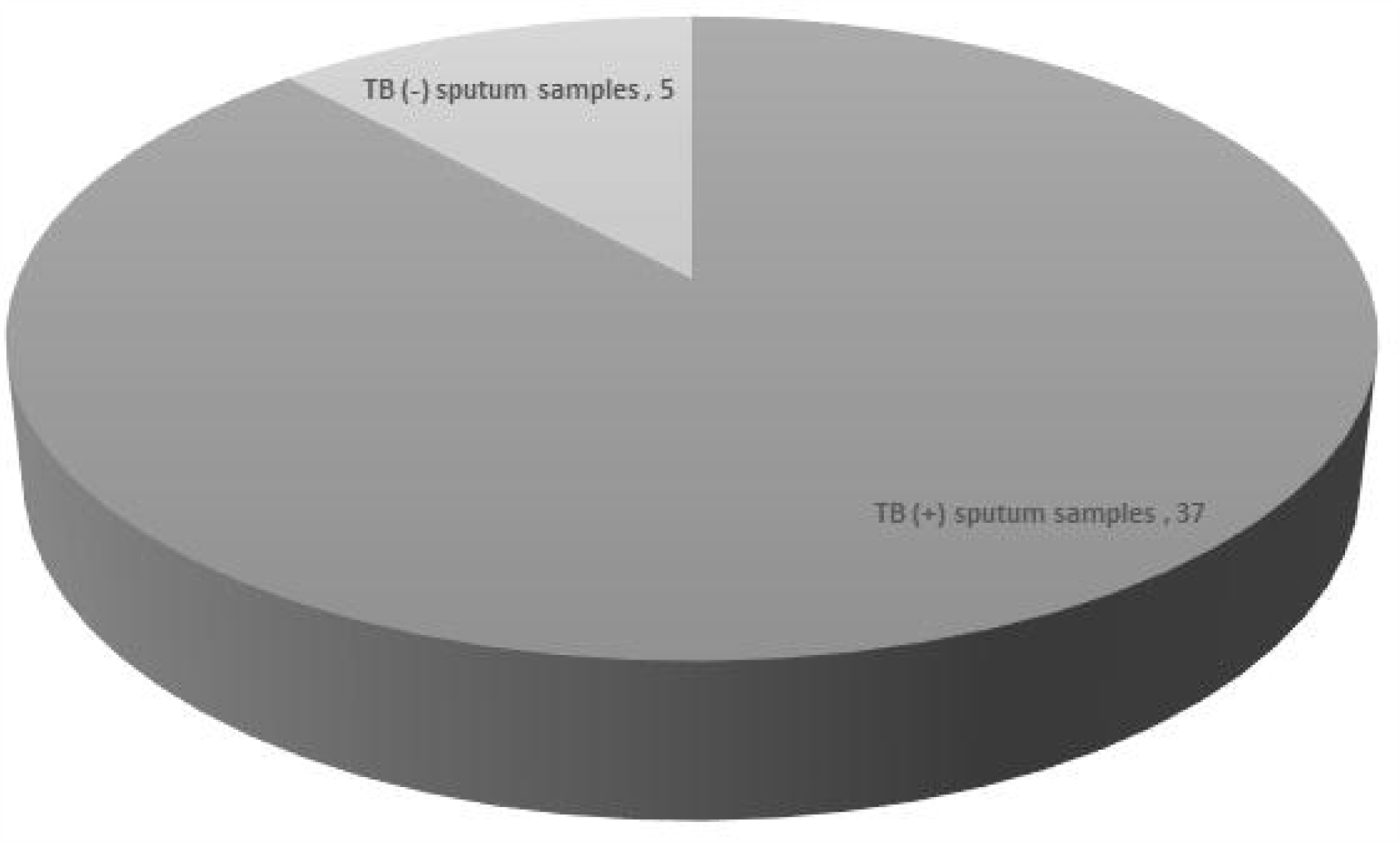
Proportion of TB-positivity in clinical samples used in this study.

Sputum samples were processed following standard bacteriological methods. Briefly, 1 ml of sputum sample was treated by adding equal volume of N-acetyl-L-cysteine–sodium hydroxide solution followed by an incubation of 20 minutes. The solution was then neutralized by adding phosphate buffered saline and centrifuged. The sediment was suspended in 1 ml volume of sterilized phosphate buffer and used for preparation of smears for microscopy as well as for inoculation into the culture media (Kent and Kubica, 1985). Liquid culture was performed using the BACTEC MGIT 960 system (Becton Dickinson, Sparks, MD). Time to culture positivity was recorded as the period in number of days between the inoculation of a sample and detection of growth of *Mycobacteria* if any.

### Loading the TBSend card with sputum

TBSend cards, routinely used for spotting processed sputum followed by its storage and extraction of DNA, were used in this study. To load the card with sputum, around 600 μL of sputum sample was processed by mixing equal volume of Spotting buffer (supplied with the kit), vortexed for 30 seconds and incubated at room temperature for 10 minutes. Around 1.2 ml of this sample-buffer solution was spotted onto the TBSend card using a disposable pipette (provided with the kit).

### Rescue of MTB bacilli from TBSend card spotted with TB infected sputum

After (a) 1 minute and (b) 15 hours from the time of spotting of the TBSend card with processed TB-positive sputum, the circular sputum-spotted card was removed from the container using the flexible handle attached to it and placed into a wide-mouth bottle prefilled with 3 ml of phosphate buffer, saline, mixed well and incubated for 15 minutes on a rotary shaker set to function at a speed of 75 revolutions per minute. Post incubation, a volume of 1 ml of the resultant “washed” buffer was transferred onto a sterile tube, processed in a way similar to that for sputum and used to run a BACTEC™ MGIT 960™ TB System for detection of live Mtb bacilli.

Each of the 42 sputum samples were divided into 3 aliquots, namely A, B and C, each comprising of 1 ml of sputum. Aliquots A and B wereprocessed andused tospot two TBSend cards labelled Card A and B respectively while aliquot C was used to inoculate growth media for running the BACTEC™ MGIT 960™ TB System.

All Card A were spotted on day 1 at 19 hours while Card B, on day 2 at 9:59 hours respectively. The process for release of probable live Mtb bacilli from Card A & B if any, was initiated at 10 hours on day 2 by washing the spotted cards as described above and used to inoculate growth tubes of the BACTEC™ MGIT 960™ TB System.

### Experimental design

For each TB positive sputum sample, three different BACTEC™ MGIT 960™ TB runs were conducted. In the first two runs, namely that for card A and B, the inoculum used was card-washed buffer retrieved after (a) 1 minute and (b) 15 hours of spotting of a TB Send card with the sputum respectively. The third run (aliquot C) was direct inoculation of sputum to run the BACTEC™ MGIT 960™ TB System after its necessary pre-inoculation processing as suggested by the manufacturer (Figure 2).

**Figure 2.**
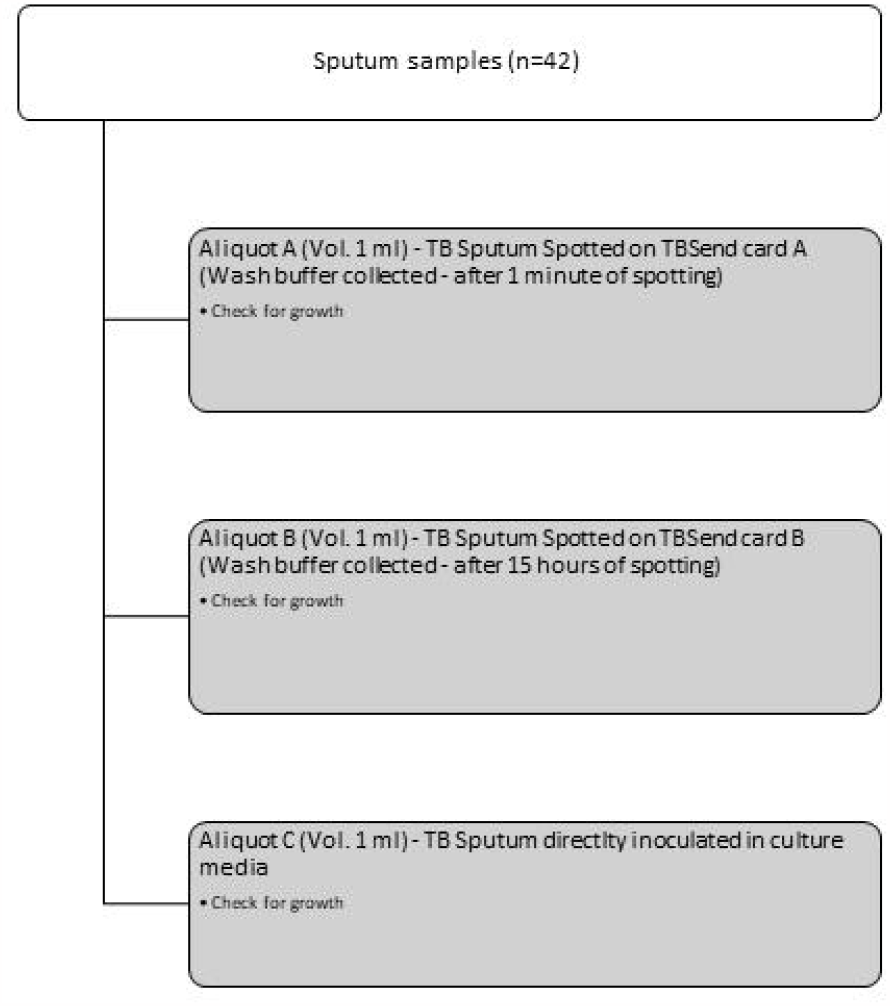
Study design for estimating infectivity of TB-sputum spotted TBSend using BACTEC^™^ MGIT 960^™^ TB System as the live TB bacilli detection platform.

Ethical approval for this study was obtained from Nirmal Hospital Private Limited Ethics Committee and with the reference number as: Nirmal/HPL/Ethics/001.

## Results

Sputum sample from 58 individuals were screened using smear microscopy technique and 37 were found to be smear positive for Mtb. Sputum samples collected from 5 healthy individuals were smear-negative as expected.

All sputum samples positive by smear microscopy were processed on the BACTEC™ MGIT 960™ TB System and found to be positive for Mtb. However, the time to culture positivity varied from sample to sample. Five sputum samples which were collected from healthy individuals did not show any growth in culture after an incubation of 42 days.

TBSend card-washed buffers taken after 1 minute and 15 hours from the time of spotting of TB positive sputum onto it did not show any Mtb growth in culture. This included all 37 sputum-positive and 5 sputum-negative samples collected from healthy individuals. The results are summarized in Table 1.

**Table 1:** Summary of data obtained from processing of the resource population that comprised of 37 *M. tuberculosis* infected sputum and 5 sputa from healthy individuals.

| Sr. no. | Sample ID | Smear Microscopy | BACTEC™ MGIT 960™ TB System |  |  |  |  |  |
| --- | --- | --- | --- | --- | --- | --- | --- | --- |
|  |  | Sputum | TBSend card A (1 minute) |  | TBSend card B (15 hours) |  | Direct Sputum |  |
|  |  | Result | Growth | Days of incubation | Growth | Days of incubation | Growth | Days of incubation |
| 1 | WM-2 | Positive | No Growth | 42 | No Growth | 42 | Growth | 27 |
| 2 | WM-3 | Positive | No Growth | 42 | No Growth | 42 | Growth | 23 |
| 3 | WM-4 | Positive | No Growth | 42 | No Growth | 42 | Growth | 18 |
| 4 | WM-26 | Positive | No Growth | 42 | No Growth | 42 | Growth | 29 |
| 5 | WM-28 | Positive | No Growth | 42 | No Growth | 42 | Growth | 24 |
| 6 | WM-29 | Positive | No Growth | 42 | No Growth | 42 | Growth | 29 |
| 7 | WM-35 | Positive | No Growth | 42 | No Growth | 42 | Growth | 27 |
| 8 | WM-40 | Positive | No Growth | 42 | No Growth | 42 | Growth | 29 |
| 9 | WM-48 | Positive | No Growth | 42 | No Growth | 42 | Growth | 28 |
| 10 | WM-49 | Positive | No Growth | 42 | No Growth | 42 | Growth | 28 |
| 11 | WM-50 | Positive | No Growth | 42 | No Growth | 42 | Growth | 32 |
| 12 | WM-57 | Positive | No Growth | 42 | No Growth | 42 | Growth | 22 |
| 13 | WM-59 | Positive | No Growth | 42 | No Growth | 42 | Growth | 25 |
| 14 | WM-60 | Positive | No Growth | 42 | No Growth | 42 | Growth | 27 |
| 15 | WM-64 | Positive | No Growth | 42 | No Growth | 42 | Growth | 28 |
| 16 | WM-109 | Positive | No Growth | 42 | No Growth | 42 | Growth | 17 |
| 17 | WM-121 | Positive | No Growth | 42 | No Growth | 42 | Growth | 26 |
| 18 | WM-124 | Positive | No Growth | 42 | No Growth | 42 | Growth | 17 |
| 19 | WM-128 | Positive | No Growth | 42 | No Growth | 42 | Growth | 21 |
| 20 | WM-129 | Positive | No Growth | 42 | No Growth | 42 | Growth | 21 |
| 21 | WM-166 | Positive | No Growth | 42 | No Growth | 42 | Growth | 11 |
| 22 | WM-181 | Positive | No Growth | 42 | No Growth | 42 | Growth | 20 |
| 23 | WM-209 | Positive | No Growth | 42 | No Growth | 42 | Growth | 38 |
| 24 | WM-213 | Positive | No Growth | 42 | No Growth | 42 | Growth | 29 |
| 25 | WM-232 | Positive | No Growth | 42 | No Growth | 42 | Growth | 36 |
| 26 | WM-313 | Positive | No Growth | 42 | No Growth | 42 | Growth | 11 |
| 27 | WM-353 | Positive | No Growth | 42 | No Growth | 42 | Growth | 21 |
| 28 | WM-354 | Positive | No Growth | 42 | No Growth | 42 | Growth | 22 |
| 29 | WM-355 | Positive | No Growth | 42 | No Growth | 42 | Growth | 22 |
| 30 | WM-356 | Positive | No Growth | 42 | No Growth | 42 | Growth | 29 |
| 31 | WM-357 | Positive | No Growth | 42 | No Growth | 42 | Growth | 35 |
| 32 | WM-358 | Positive | No Growth | 42 | No Growth | 42 | Growth | 18 |
| 33 | WM-360 | Positive | No Growth | 42 | No Growth | 42 | Growth | 25 |
| 34 | WM-361 | Positive | No Growth | 42 | No Growth | 42 | Growth | 8 |
| 35 | WM-362 | Positive | No Growth | 42 | No Growth | 42 | Growth | 18 |
| 36 | WM-363 | Positive | No Growth | 42 | No Growth | 42 | Growth | 25 |
| 37 | WM-368 | Positive | No Growth | 42 | No Growth | 42 | Growth | 31 |
| 38 | WM-24 | Negative | No Growth | 42 | No Growth | 42 | No Growth | 42 |
| 39 | WM-85 | Negative | No Growth | 42 | No Growth | 42 | No Growth | 42 |
| 40 | WM-107 | Negative | No Growth | 42 | No Growth | 42 | No Growth | 42 |
| 41 | WM-110 | Negative | No Growth | 42 | No Growth | 42 | No Growth | 42 |
| 42 | WM-125 | Negative | No Growth | 42 | No Growth | 42 | No Growth | 42 |

## Discussion

In the domain of infectious diseases that are transmitted by air, Mtb is a classical archetype and an extremely potent causative agent for the disease (Roy and Milton, 2004). In the case of human tuberculosis, the primary source of infection is other humans suffering from this pulmonary disease. Zoonotic tuberculosis, mainly arising out of cattle with *M. bovis* being the pathogen of cause do carry significance but its contribution to the cause of human tuberculosis is low and at 1.4% only (Müller *et al*., 2013). The diagnosis of tuberculosis classically depends on detection of Mtb pathogen by culture method. Acid fast bacilli or AFB smear is an economical, inexpensive method for TB diagnosis but cannot differentiate nontuberculous mycobacteria (NTM) from Mtb. The culture method of detection of Mtb is more sensitive compared to AFB smear method but the results are generated after several weeks making it a time-consuming protocol. (Somoskövi *et al*., 2000). Compared to these two methods, Nucleic acid amplification tests or NAATs has now evolved as a rapid and sensitive technology for diagnosis of TB and is also effective in distinguishing NTMs (Su *et al*., 2000). A large number of studies have highlighted the advantages of NAAT in controlling TB which primarily includes avoiding unnecessary treatment of the disease (Bourgi *et al*., 2017; Marks *et al*., 2013), shorter time period and reduced delays in initiation of treatment (Peralta *et al*., 2016; Singla *et al*., 2014).

In India, observation suggests that detection of drug resistant TB, which was earlier dominated by the culture method has now gradually shifted towards molecular drug susceptibility testing (Molecular-DST). Several NAAT protocols has been approved by the World Health Organization (WHO) which include line probe assays (LPA), namely GenoType® MTBDRplus VER 2.0 (first-line LPA) and GenoType® MTBDRsl VER 2.0 (second line LPA), manufactured by Hain Lifesciences, Nehren, Germany, Xpert® MTB/RIF & MTB/RIF Ultra assay (Cepheid, Sunnyvale, USA) and Truenat™ MTB-Rif Dx assay (Molbio Diagnostics, Goa, India) (World Health Organization, 2020). The Line Probe Assay from Hain Lifesciences has been adopted In India by the NTEP programme and is the platform of choice for detecting drug resistant TB in direct smear-positive sputum samples and cultured Mtb bacteria obtained from smear negative samples (RNTCP, 2017). Data suggest that in India, in the year 2018, around 3,46,282 and 72,748 samples were tested using first- and second-line LPA as compared to only 16,399 samples tested by culture method for detection of drug resistant TB (NTEP, 2020). This hints at the scalability of adopting NAAT for detection of drug resistant tuberculosis with the LPA testing platform serving as a model. However, it may be noted that the use of LPA is limited to the national and intermediate reference laboratories and few certified laboratories in the country which are equipped with high end diagnostic facilities and adequately trained operators that are not present in Designated Microscopy Centres (NTEP, 2020). This gap raises potential logistic challenge as it involves transportation of highly infectious sputum from remote and geographically challenging areas to the central testing laboratories and poses potential biosafety concerns.

TBSend card, developed by Wobble Base Bioresearch Private limited, India (Patent Application number: 201621024943 dated July 20, 2016) is a proprietary reagent coated cellulose matrix housed in an air tight container that can be used for spotting Mtb infected sputum. For applying sputum on the card, it is mixed with an equal volume of spotting buffer at a ratio of 1:1 and the resultant mixture is poured onto the card. Mtb-infected sputum-spotted cards retain the target pathogen DNA for a long period of time (1 day to > 6 years) and can release it on demand by simply suspending it in a buffer and incubating for 15 minutes before running a CB-NAAT. It can also be purified for other NAAT which are based on classical real time fluorescent PCR, end point PCR, Line probe assays and Next generation sequencing platforms, using a hybrid spin column (Patent application No. 201921017045 dated April 29, 2019) that is specially designed for this purpose and similar other DNA purification applications.

Some of the crucial components of TBSend card module are the spotting buffer and the reagents used to coat the card which comprises of strong chaotropic agents and nonionic surfactants having hydrophilic polyethylene oxide chain and aromatic hydrocarbon groups with potential to lyse cells and also an anti-fungal agent. Chaotropic salts used in this module are co-solutes that break down the network of hydrogen bonds between the water molecules and effectively lower down the stability of biological proteins encountered in the sputum sample by reducing the hydrophobic effect (Salvi *et al*., 2005).

Most of the commercial nucleic acid extraction kits are known to contain strong chaotropic salts. They are more effective than traditional cell lysis buffers that comprises of NaCl, EDTA and Tris buffer as the primary components (Eslami *et al*., 2017).

In a study by Clinghan *et al*., (2013) the authors demonstrated that two popular DNA extraction kits, namely NucliSENS easyMAG (bioMérieux, Boxtel, the Netherlands) and Qiagen QIAamp DNA mini kit (Hilden, Germany) effectively inactivated Mtb cells in clinical samples rendering them biocontainment-safe for handling purposes. For this, clinicals strains of Mtb were incubated in NucliSENS and Qiagen lysis buffer, both of which contain chaotropic salt as the primary cell lysing agent and then inoculated onto LJ media contained slant-tubes and MGIT growth tubes. No growth was observed after 6 weeks in either of the samples.

In this study, the resource population was carefully constructed in order to involve a majority number of TB positive sputum samples in the study. All sputum samples found positive by smear microscopy also demonstrated growth in the BACTEC™ MGIT 960™ TB System. As expected, the days to positivity from the date of inoculation for each sample varied depending upon the quantum of live inoculum that went into the growth tubes for each of the samples.

TBSend card spotted sputum cards washed with phosphate buffer by incubating for 15 minutes in a rotary shaker did not release any live bacilli. This was demonstrated by nil growth in culture (when card-washed phosphate buffers were used as inoculum) after an incubation of 42 days, the maximum recommended by the manufacturer of BACTEC™ MGIT 960™ TB System to call a sample as TB positive.

Two categories of TBSend cards were used in this study. In the first (Card A), the TB-sputum spotted cards were washed in phosphate buffer 1 minute after spotting while in the second (Card B), it was washed after 15 hours after spotting with TB-infected sputum. In the former, the time lapse after the sputum sample came in contact with the card was less (1 minute) compared to the other (card B) where the card remained in contact with the sputum for up to a period of 15 hours. In both the cases no live Mtb microbes were detected by culture method indicating that Mtb cell viability was lost at least 1 minute after deposition of the processed samples onto the card if not earlier.

The complete absence of any live Mtb on the TBSend cards spotted with processed TB positive sputum is well anticipated. The spotting buffer to which sputum samples were mixed at a ratio of 1:1 comprised of a mixture of strong chaotropic salts among other components, all in soluble form. On the other hand, the cellulose matrix with which the processed sputum (mixed with spotting buffer) came in contact during the spotting process too had chaotropic salts and non-ionic detergents that played crucial role in lysing of cells. This study therefore further consolidated the finding of Clinghan *et al*., (2013) and established the fact that chaotropic agents effectively inactivated live Mtb bacilli.

From this study, the authors concluded that once Mtb infected sputum is spotted on to a TBSend card following prescribed instructions, it achieved optimal biosafety quotient and could be labelled as biosafe and non-hazardous from Mtb infection.

## Acknowledgements

This study was supported by Grand Challenge-TB Control (GCTBC) grants, namely GCTBC/C2P1/2015/09/30/01 & GCTBC/C2P2/2017/04/01/01 and funded by Industry Research Assistance Council (BIRAC) of the Government of India, United States Agency for International Development (USAID), and Bill & Melinda Gates Foundation. IKP Knowledge Park (Hyderabad, India) was the implementation partner for these programmes.

## References

Basu, M., Sinha, D., Das, P., Roy, B., Biswas, S., & Chattopadhyay, S. (2013). Knowledgeandpracticeregardingpulmonary tuberculosis among private practitioners. Indian Journal of Community Health, 25(4), 403–412.

Bhatt, G., Vyas, S.,& Trivedil, K. (2012). Anepidemiologicalstudyof multi drug resistant tuberculosis cases registered under Revised National Tuberculosis Control Programme of Ahmedabad City. The Indian journal of tuberculosis, 59(1), 18–27.

Bourgi, K., Patel, J., Samuel, L., Kieca, A., Johnson, L., & Alangaden, G. (2017). Clinical Impact of Nucleic Acid Amplification Testing in the Diagnosis of Mycobacterium Tuberculosis: A 10-Year Longitudinal Study. Open forum infectious diseases, 4(2), ofx045.

Clinghan, R., Anderson, T. P., Everett, V., & Murdoch, D. R. (2013). Viability of Mycobacterium tuberculosis after processing with commercial nucleic acid extraction kits. Journal of clinical microbiology, 51(6), 2010.

Eslami, G., Khalatbari-Limaki, S., Ehrampoush, M. H., Gholamrezaei, M., Hajimohammadi, B., & Oryan, A. (2017). Comparison of Three Different DNA Extraction Methods for Linguatula serrata as a Food Born Pathogen. Iranian journal of parasitology, 12(2), 236–242.

Galvani, A. P., & May, R. M. (2005). Epidemiology: dimensions of superspreading. Nature, 438(7066), 293–295.

Jangid VK, Agrawal NK, Yadav GS, Pandey S, Mathur BB. (2016). Health-seeking behaviour and social stigma for tuberculosis in tuberculosis patients at a tertiary care center in North West India. Int J Med Sci Public Health, 5(9):1893–99.

Kent PT, Kubica GP (1985) Public health microbiology, a guide for the level III laboratory. Centres for Disease Control, Division of Laboratory Training and Consultation, Atlanta, GA.

Kramer, A., Schwebke, I., & Kampf, G. (2006). How long do nosocomial pathogens persist on inanimate surfaces? A systematic review. BMC infectious diseases, 6, 130.

Marks, S. M., Cronin, W., Venkatappa, T., Maltas, G., Chon, S., Sharnprapai, S., Gaeddert, M., Tapia, J., Dorman, S. E., Etkind, S., Crosby, C., Blumberg, H. M., & Bernardo, J. (2013). The healthsystem benefits and cost-effectiveness of using Mycobacterium tuberculosis direct nucleic acid amplification testing to diagnose tuberculosis disease in the United States. Clinical infectious diseases: an official publication of the Infectious Diseases Society of America, 57(4), 532–542.

Müller, B., Dürr, S., Alonso, S., Hattendorf, J., Laisse, C. J., Parsons, S. D., van Helden, P. D., & Zinsstag, J. (2013). Zoonotic Mycobacterium bovis-induced tuberculosis in humans. Emerging infectious diseases, 19(6), 899–908.

NTEP. (2020). India TB Report-National Tuberculosis Elimination Programme Annual Report. India, National Tuberculosis Elimination Programme.

Peralta, G., Barry, P., & Pascopella, L. (2016). Use of Nucleic Acid Amplification Tests in Tuberculosis Patients in California, 2010-2013. Open forum infectious diseases, 3(4), ofw230.

Phillips, C. J., Foster, C. R., Morris, P. A., & Teverson, R. (2003). The transmission of Mycobacterium bovis infection to cattle. Research in veterinary science, 74(1), 1–15.

Rekha, T., Singh, P., Unnikrishnan, B., Prasanna Mithra, P., Kumar, N., Prasad, K. D., Raina, V., Kumar Papanna, M., & Kulkarni, V. (2013). Sputum collection and disposal among pulmonary tuberculosis patients in coastal South India. The international journal of tuberculosis and lung disease: the official journal of the International Union against Tuberculosis and Lung Disease, 17(5), 621–623.

Revised National Tuberculosis Control Programme (RNTCP). (2017). Guidelines on Programmatic Management of Drug Resistant TB (PMDT) in India. India, Revised National TB Control Program.

Roy, C. J., & Milton, D. K. (2004). Airborne transmission of communicable infection--the elusive pathway. The New England journal of medicine, 350(17), 1710–1712.

Salvi, G., De Los Rios, P., & Vendruscolo, M. (2005). Effective interactions between chaotropic agents and proteins. Proteins, 61(3), 492–499.

Singla, N., Satyanarayana, S., Sachdeva, K. S., Van den Bergh, R., Reid, T., Tayler-Smith, K., Myneedu, V. P., Ali, E., Enarson, D. A., Behera, D., & Sarin, R. (2014). Impact of introducing the line probe assay on time to treatment initiation of MDR-TB in Delhi, India. PloS one, 9(7), e102989.

Somoskövi, A., Ködmön, C., Lantos, A., Bártfai, Z., Tamási, L., Füzy, J., & Magyar, P. (2000). Comparison of recoveries of mycobacterium tuberculosis using the automated BACTEC MGIT 960 system, the BACTEC 460 TB system, and Löwenstein-Jensen medium. Journal of clinical microbiology, 38(6), 2395–2397.

Su WJ, Tsou AP, Yang MH, Huang CY, Perng RP. Clinical experience in using polymerase chain reaction for rapid diagnosis of pulmonary tuberculosis. Zhonghua Yi Xue Za Zhi (Taipei). 2000;63(7):521–6.

TBC India. (2013). Diagnosis of smear-positive pulmonary TB. New guidelines. New Delhi, India. Available from URL: http://www.tbindia.nic.in/pdf/RNTCPlabNetworkGuidelines (Accessed November 2013).

Urdahl, K. B., Shafiani, S., & Ernst, J. D. (2011). Initiation and regulation of T-cell responses in tuberculosis. Mucosal immunology, 4(3), 288–293.

Wang, C. S., Chen, H. C., Chong, I. W., Hwang, J. J., & Huang, M. S. (2008). Predictors for identifying the most infectious pulmonary tuberculosis patient. Journal of the Formosan Medical Association = Taiwan yi zhi, 107(1), 13–20.

Wolf, A. J., Desvignes, L., Linas, B., Banaiee, N., Tamura, T., Takatsu, K.,& Ernst, J. D.(2008). Initiationof theadaptiveimmuneresponse to Mycobacterium tuberculosis depends on antigen production in the local lymph node, not the lungs. The Journal of experimental medicine, 205(1), 105–115.

Woolhouse, M. E., Dye, C., Etard, J. F., Smith, T., Charlwood, J. D., Garnett, G. P., Hagan, P., Hii, J. L., Ndhlovu, P. D., Quinnell, R. J., Watts, C. H., Chandiwana, S. K., & Anderson, R. M. (1997). Heterogeneities in the transmission of infectious agents: implicationsforthedesignofcontrolprograms. Proceedingsofthe National Academy of Sciences of the United States of America, 94(1), 338–342.

World Health Organization (2015), A Global Action Framework for TB Research in Support of the Third Pillar of WHO’s end TB Strategy. Geneva. WHO/HTM/TB/2015.26.

World Health Organization. (2007). Blood safety and clinical technology: guideline on standard operating procedures on microbiology. Chapter 17: tuberculosis. Geneva, Switzerland: WHO, 2007. Available from URL: http://209.61.208.233/en/Section10/Section17/Section53/Section482.htm [Accessed November 2013].

World Health Organization. (2020). WHO consolidated guidelines on tuberculosis: module 3: diagnosis: rapid diagnostics for tuberculosis detection. World Health Organization. https://apps.who.int/iris/handle/10665/332862. License: CC BY-NC-SA 3.0 IGO

Yang, S., Lee, G. W., Chen, C. M., Wu, C. C., & Yu, K. P. (2007). The size and concentration of droplets generated by coughing in human subjects. Journalof aerosol medicine: the official journal of the International Society for Aerosols in Medicine, 20(4), 484–494.

